# Ambient noise from ocean surf drives frequency shifts in non-passerine bird song

**DOI:** 10.1101/2020.06.17.156232

**Authors:** Matteo Sebastianelli, Daniel T. Blumstein, Alexander N. G. Kirschel

## Abstract

Effective communication in birds is often hampered by background noise, with many recent studies focusing on the effect of anthropogenic noise on passerine bird song. Continuous low-frequency natural noise is predicted to drive changes in both frequency and temporal patterning of bird vocalizations, but the extent to which these effects may also affect birds that lack vocal learning is not yet fully understood. Here we use a gradient of exposure to natural low-frequency noise to assess whether it exerts selective pressure on vocalizations in a species whose songs are innate. We tested whether three species of *Pogoniulus* tinkerbirds adapt their song when exposed to a source of continuous low-frequency noise from ocean surf. We show that dominant frequency increases the closer birds are to the coast in all the three species, and in line with higher noise levels, indicating that ocean surf sound may apply a selective pressure on tinkerbird songs. As a consequence, tinkerbirds adapt their songs with an increase in frequency to avoid the masking effect due to overlapping frequencies with ambient noise, therefore improving long-range communication with intended receivers. Our study provides for the first time, compelling evidence that natural ambient noise affects vocalizations in birds whose songs are developed innately. We believe that our results can also be extrapolated in the context of anthropogenic noise pollution, hence providing a baseline for the study of the effects of low-frequency ambient noise on birds that lack vocal learning.

**Significance Statement:** Birdsong is constantly under selection as it mediates key interactions such as mate attraction, competition with same-sex individuals for reproduction and competition with heterospecifics for space-related resources. Any phenomenon that interferes with communication can therefore have a profound impact on individual fitness. Passerines are more likely to avoid the masking effect of background noise because of their higher vocal flexibility. Many non-passerine species lacking such flexibility might therefore be more vulnerable to the negative effects on their fitness of exposure to low-frequency background noise. Species incapable of adapting their signals to background noise are predicted to disappear from noisy areas. Despite this, we show that species that lack song learning may show an adaptive response to natural noise which may develop over evolutionary timescales.

## Introduction

Many species rely on acoustic communication to accomplish functions that are crucial for their survival (Bradbury and Vehrencamp 2011). Bird song, for instance, has been shown to mediate behaviours involved in mate attraction, competition for partners, food and space (Catchpole and Slater 2008; Naguib and Riebel 2014); even though it may also function to coordinate group movements and to warn other individuals against potential threats (Naguib and Wiley 2001; Bradbury and Vehrencamp 2011; Halfwerk et al. 2018). An effective signal transfer is therefore essential to ensure the prompt behavioural response of the receiver.

The transfer of clear signals might be hampered by the sound transmission properties of the environment, which may degrade signals (Brumm and Naguib 2009), or by interference from environmental noise (Brumm and Slabbekoorn 2005; Blumstein et al. 2011). Under the latter scenario, sounds similar in frequency and amplitude can have a masking effect and potentially lead to the transmission of incomplete or incorrect information (Slabbekoorn 2013). Such effects have a strong effect on vocal behaviour of birds (Patricelli and Blickley 2006; Slabbekoorn 2013). Indeed, experiments have shown birds in one environment with a specific ambient noise profile respond less to songs adapted to different ambient noise profiles than to those adapted to similar ambient noise profiles (Kirschel et al. 2011). Therefore, loud and continuous background noise impose strong selective pressures on bird song to increase its effectiveness in noisy environments (Slabbekoorn and Smith 2002; Brumm and Slabbekoorn 2005; Patricelli and Blickley 2006; Slabbekoorn and den Boer-Visser 2006; Slabbekoorn and Ripmeester 2008; Halfwerk and Slabbekoorn 2009; Nemeth and Brumm 2010).

Birds react to low frequency ambient noise pressure in different ways (Brumm and Slabbekoorn 2005; Swaddle et al. 2015). Some have been shown to increase their minimum frequency (Slabbekoorn and den Boer-Visser 2006; Nemeth and Brumm 2009, 2010; Hu and Cardoso 2010; Mendes et al. 2011; Ríos-Chelén et al. 2012), maximum frequency (Francis et al. 2011; Mendes et al. 2011) and others their dominant frequency (Nemeth and Brumm 2009; Hu and Cardoso 2010; Proppe et al. 2011, 2012; Lazerte et al. 2016; Luther et al. 2016; LaZerte et al. 2017; Tolentino et al. 2018) in response to background noise. Increases in frequency may, however, be a side effect of singing at higher amplitude in noisy environments (Nemeth and Brumm 2010) – the Lombard Effect (Brumm and Zollinger 2011; Zollinger and Brumm 2011) – as amplitude and song frequency are often correlated (Beckers et al. 2003; Amador et al. 2008; Zollinger et al. 2012). Other adaptations to low-frequency ambient noise include increasing signal redundancy (Brumm and Slater 2006; Deoniziak and Osiejuk 2016), singing more often (Deoniziak and Osiejuk 2019), for longer periods (Brumm and Slater 2006; Nemeth and Brumm 2009; Sierro et al. 2017) or at specific time intervals (Dominoni et al. 2016).

Changes in vocal parameters can result from different mechanisms, for instance, response to background noise might be plastic, as found in House Finches (*Carpodacus mexicanus*) (Bermúdez-Cuamatzin et al. 2009), or learned, as demonstrated in Black-capped chickadees (*Poecile atricapillus*) (Lazerte et al. 2016) and White-crowned sparrows (*Zonotrichia laucophyrs*) (Moseley et al. 2018). Shifts in signal design might also arise because selection may favor individuals that minimize the masking effect of ambient noise (Slabbekoorn and Smith 2002; Kirschel et al. 2009a, 2011). This scenario is compatible with the song developing by sensory drive (Endler 1992), a mechanism which appears to have shaped acoustic signals of many Neotropical suboscines (Seddon 2005) that lack song learning capabilities (Touchton et al. 2014).

Most studies on the effects of noise on acoustic communication have addressed this issue by looking at the effects of anthropogenic noise pollution. However, natural sources of noise may have similar masking effects on animal signalling (Davidson et al. 2017; Goutte et al. 2018). For instance, Halfwerk et al (2016) show multimodal communication between male Tungara frogs (*Physalaemus pustulosus*) was hindered when geophonic noise from windy and rainy conditions was simulated. Other studies on birds and other taxa have also shown an effect of natural background noise on communication (Lengagne et al. 1999; Lengagne and Slater 2002; Brumm and Slater 2006; Feng et al. 2006; Kirschel et al. 2009a; Davidson et al. 2017). Therefore, natural ambient noise is likely to be as impactful as anthropogenic noise and with such noise present over evolutionary timescales it is likely to have evolutionary implications for acoustic communication (Davidson et al. 2017).

To date, the study of the effects of ambient noise on bird signalling has focused mostly on oscine passerines that learn their songs by way of auditory feedback (Hu and Cardoso 2010; Ríos-Chelén et al. 2012). By contrast, there is scant information on how taxa that lack vocal learning, such as suboscines and many non-passerines birds, cope with high background noise levels (Gentry et al. 2018; Tolentino et al. 2018). Studies on non-passerines include those on King penguins (*Apten odytes*) (Lengagne et al. 1999) and Tawny owls (Lengagne and Slater 2002). In both cases, responses to increased ambient noise were in temporal patterning of their vocalizations. King penguins increased both the number of calls and syllables per call emitted under strong winds, whereas Tawny owls reduced call rates under rainy conditions because the interference of rain noise increased the unreliability of the information conveyed in their calls. Hu and Cardoso (2010) did document changes in the frequency domain in response to anthropogenic noise in a non-passerine by observing an increase in minimum frequency in urban rainbow lorikeets (*Tricoglossus haematodus*) and eastern rosellas (*Platycercus eximius*) two Psittaculidae (Order: Psittaciformes). However, parrots, like hummingbirds and oscine passerines, are capable of vocal learning (Nottebohm 1972; Kroodsma 1982; Saranathan et al. 2007; Catchpole and Slater 2008) and therefore may respond plastically to increased background noise levels (Osmanski and Dooling 2009; Scarl and Bradbury 2009). Although birds not capable of learning such as suboscines and many non-passerines may be more vulnerable to the effects of increased background noise given their inability to adapt their signals (Ríos-Chelén et al. 2012), little is known about the mechanisms that ensure efficient communication under noisy conditions in such taxa.

Here, we investigate whether *Pogoniulus* tinkerbirds (Family: Lybiidae; Order: Piciformes) might adapt the frequency of their songs in response to increased geophonic ambient noise from ocean surf. Tinkerbirds emit a simple, single pitch, stereotyped song that develops innately (Kirschel et al. 2009a, 2020; Nwankwo et al. 2018). Because of the absence of auditory feedback in song development, adaptation to noisy environments is unlikely to involve a learned or plastic response. Instead, any variation in tinkerbird song that would minimize the masking effect of noise may reflect an adaptive change. Hence, our study specifically addresses whether there could be a selective pressure on tinkerbird song of low frequency surf sound by focusing on species whose songs are innately developed. Previous work has found evidence for character displacement in tinkerbird song frequency when two species coexist at high densities, consistent with a role of competitive or reproductive interference of songs of similar frequencies (Kirschel et al. 2009b, 2020). We test whether yellow-throated (*Pogoniulus subsulphureus*), red-fronted (*P. pusillus*) and the coastal subspecies of yellow-rumped tinkerbird (*P. bilineatus fischeri*) adjust their song along a gradient of exposure to low-frequency ambient noise emanating from ocean surf in their coastal populations. In the case of *P. subsulphureus*, we also measure local ambient noise to test for a gradient in noise levels with distance and whether there is a direct relationship of low frequency surf sound and song frequency.

## Methods

### Study Species

*Pogoniulus* tinkerbirds are barbets (Family: Lybiidae) that are widely distributed throughout Sub-Saharan Africa. They are mostly frugivorous, feeding mainly on mistletoe, even though they also take small invertebrates (Godschalk 1985; Dowsett-Lemaire 1988; Short and Horne 2001). *P. subsulphureus* (hereafter *subsulphureus*) strictly inhabits tropical lowland rainforests in Central and Western Africa (Short and Horne 2002; Kirschel et al. 2020), whereas *P. pusillus* (hereafter *pusillus*) occupy savanna woodland and secondary forest below 2000 meters. On the other hand, *P. b. fischeri* (hereafter *fischeri*) only occurs in coastal forests in southern Kenya and on the island of Zanzibar (Nwankwo et al. 2018).

### Song Collection and Acoustical Analysis

We obtained recordings of *P. subsulphureus, P. pusillus* and *P. b. fischeri* from a total of 15 coastal locations in Cameroon and Kenya within 4 km from the shore (Fig.1). Fifty ambient noise recordings were obtained from four locations in Cameroon by taking 1-minute long recordings every hour from 7:00 to 12:00, holding the microphone horizontally every 10 seconds in each of the four cardinal direction (North, South, East, West) and then vertically upwards, as described in Kirschel et al. (2009a). Ambient noise and *subsulphureus* songs were recorded using a Marantz PMD670 a Sennheiser ME67, while *pusillus* and *fischeri* songs were recorded with a Marantz PMD661 recorder with a MKH8050 or MKH8020 microphone, the latter housed in a Telinga parabolic reflector.

**Fig. 1.**
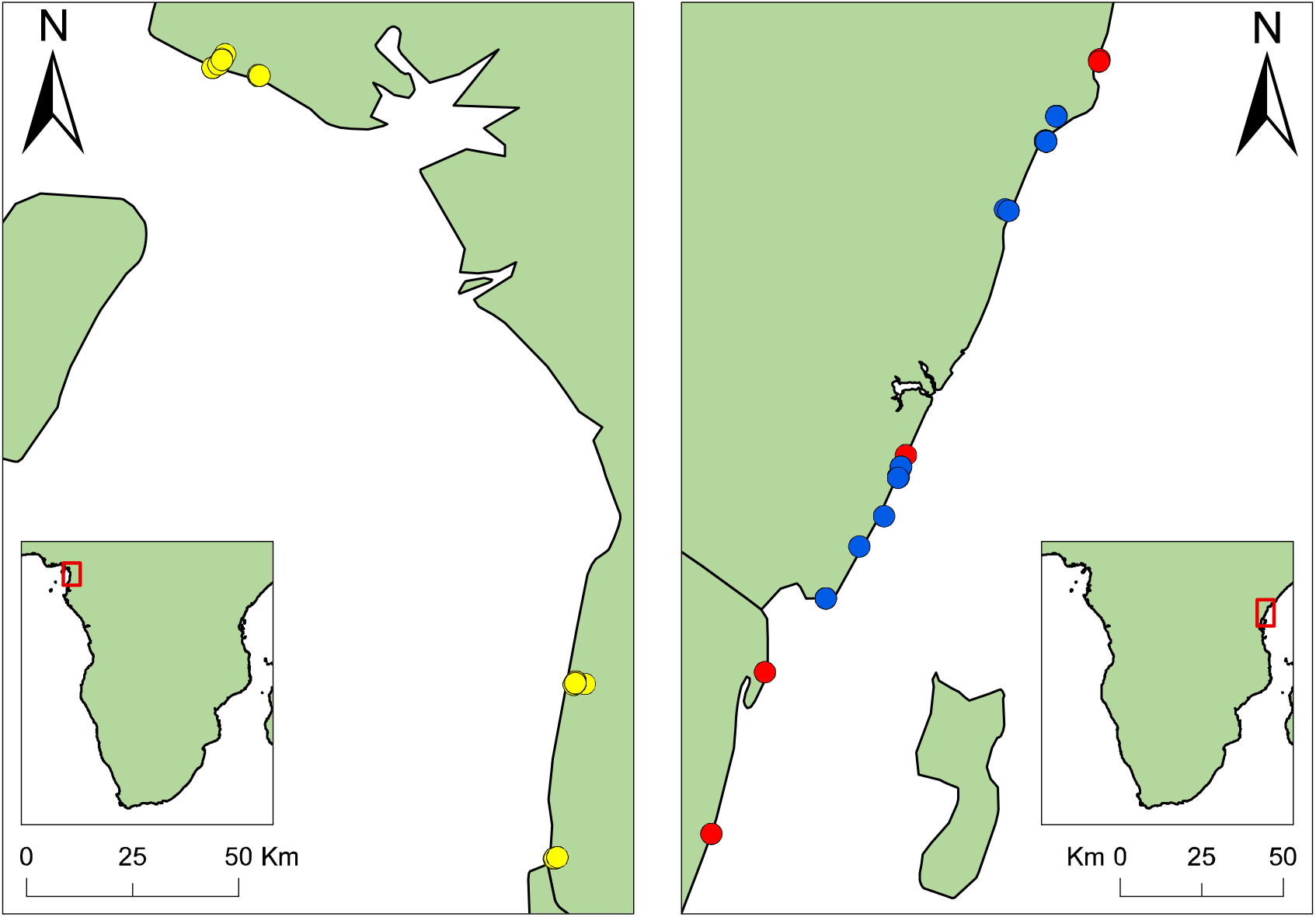
Map of recording localities in Cameroon (left) and Kenya (right). Coloured points represent recording locations of *P. subsulphureus* (yellow), *P. pusillus* (red), and *P. b. fischeri* (blue).

Recordings were saved as WAV or MP3 files and imported into Raven Pro 1.6 (Center for Conservation Bioacoustics 2019), in which songs were measured using its built-in automated energy detectors. Target signal parameters were set as follow: minimum and maximum frequencies spanned from 800 to 1700 Hz according to the species, minimum duration was 0.03 s and maximum duration 0.3; minimum separation was set to 0.01 s for *fischeri*, 0.05 s for *subsulphureus* and 0.25 for *pusillus*. The signal to noise ratio (SNR) threshold was set in order to detect the maximum number of notes and varied depending on the background noise levels on the recording. Most of the detections were obtained setting the SNR above the threshold of 10-20 dB. We chose this instead of a manual-measurement approach since the latter can lead to biased measurements (Brumm et al. 2017; Ríos-Chelén et al. 2017). Raven provided peak frequency measurements from the spectrogram view (DFT size: 4096; Window: Hanning, 3 dB; overlap: 50) and we obtained the dominant frequency by calculating the mean from peak frequency values of all notes detected on each recording.

From the 50 ambient noise recordings, six were removed because of loud anthropogenic traffic noise in the background and another was excluded because of loud stream waterfall noise, both of obscured natural surf sound. From the remaining recordings, we selected and merged together five high quality 5 s intervals per direction. In one instance we included just the four 5 s intervals from cardinal directions, because the vertical recording was beset by mechanical interference. Each 25 s song cut was then imported into R and the ambient noise amplitude (dBA) at 1 kHz was calculated using the noise profile function provided in the baRulho R package (Araya-Salas 2020). Subsequently, we used amplitude at 1 kHz (a measure of low frequency noise) as covariate in statistical models.

It was not possible to record data blind because our study was specifically focused on tinkerbirds. While *subsulphureus* in Cameroon was sampled with this specific question in mind, sampling of *pusillus* and *fischeri* was performed as part of parallel studies on song variation (e.g., Nwankwo et al. 2018).

### Spatial Distance Calculation

GPS coordinates of singing tinkerbirds and ambient noise recorded in the field were obtained using a Garmin GPSMap. We imported the coordinates into Google Earth Pro and calculated the closest distance from each recording location to the coastline using its built-in measuring tool.

### Statistical Analysis

To test whether ocean surf sound affects tinkerbird song, we measured the effect of distance from the coast on dominant frequency of *subsulphureus*, *pusillus* and *fischeri* songs. This effect was measured within 4 km from the coast as ambient noise recordings were collected within that range and songs of birds further from the coast are likely influenced by other factors, including elevation (Kirschel et al. 2009b). We assumed that, if ocean surf sound has an effect on their song, dominant frequency would decrease as the distance from the coast increases. For the coastal population of *subsulphureus* in Cameroon, for which ambient noise recordings were also available, we tested whether dominant frequency increases with background noise amplitude measured at 1 kHz, and also whether ambient noise (1 kHz) also decreases with increased distance from the coast.

We fitted Gaussian generalized linear mixed models (GLMMs) in the glmmTMB R package (Brooks et al. 2017) using log-transformed dominant frequency of *subsulphureus, pusillus* and *fischeri* as response variables in three separate models and including log-distance from the shore as a fixed factor. Bird ID nested in location were used as random factors to account for individual variation as well as variation among field sites. In the *subsulphureus* model, we also added ambient noise amplitude (measured at 1 kHz) of the closest ambient noise recording as fixed factor. We then measured the effect of distance from the coast (log-transformed) on ambient noise amplitude (1 kHz) in Cameroon coastal sites using the latter as response variable and location as random effect. *subsulphureus* models were selected according to the lowest corrected Akaike Information Criterion score. Assumptions of all models were validated using the functions provided in DHARMa (Harting 2019).

## Results

We obtained 86 recordings (39 *subsulphureus*, 21 *pusillus* and 26 *fischeri*) from a total of 65 individuals (31 *subsulphureus*, 16 *pusillus* and 18 *fischeri*) in our coastal sites in Cameroon and Kenya (Fig.1) within 4 km. Of these, 2 were sourced from Xenocanto (https://xeno-canto.org), respectively 1 for *pusillus* and 1 for *fischeri*. We found a significant negative effect of distance from the coast (within 4 km) on dominant frequency (log-transformed) in *subsulphureus, pusillus* and in *fischeri* (Fig 2, Table 1). *subsulphureus* model with both area distance from the coast and ambient noise (1kHz) was not selected because presented high AICc scores (Table S1).

**Table 1.**
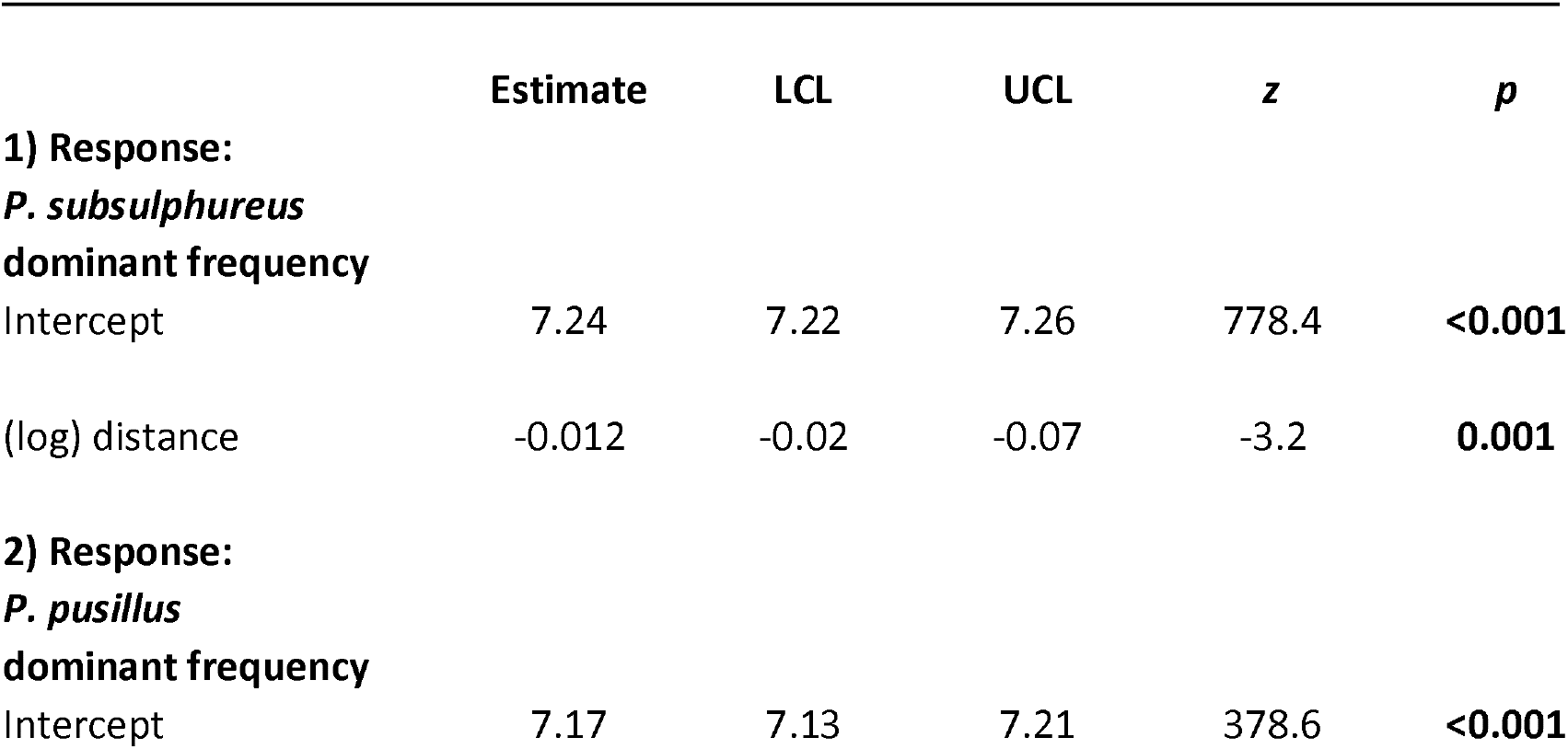
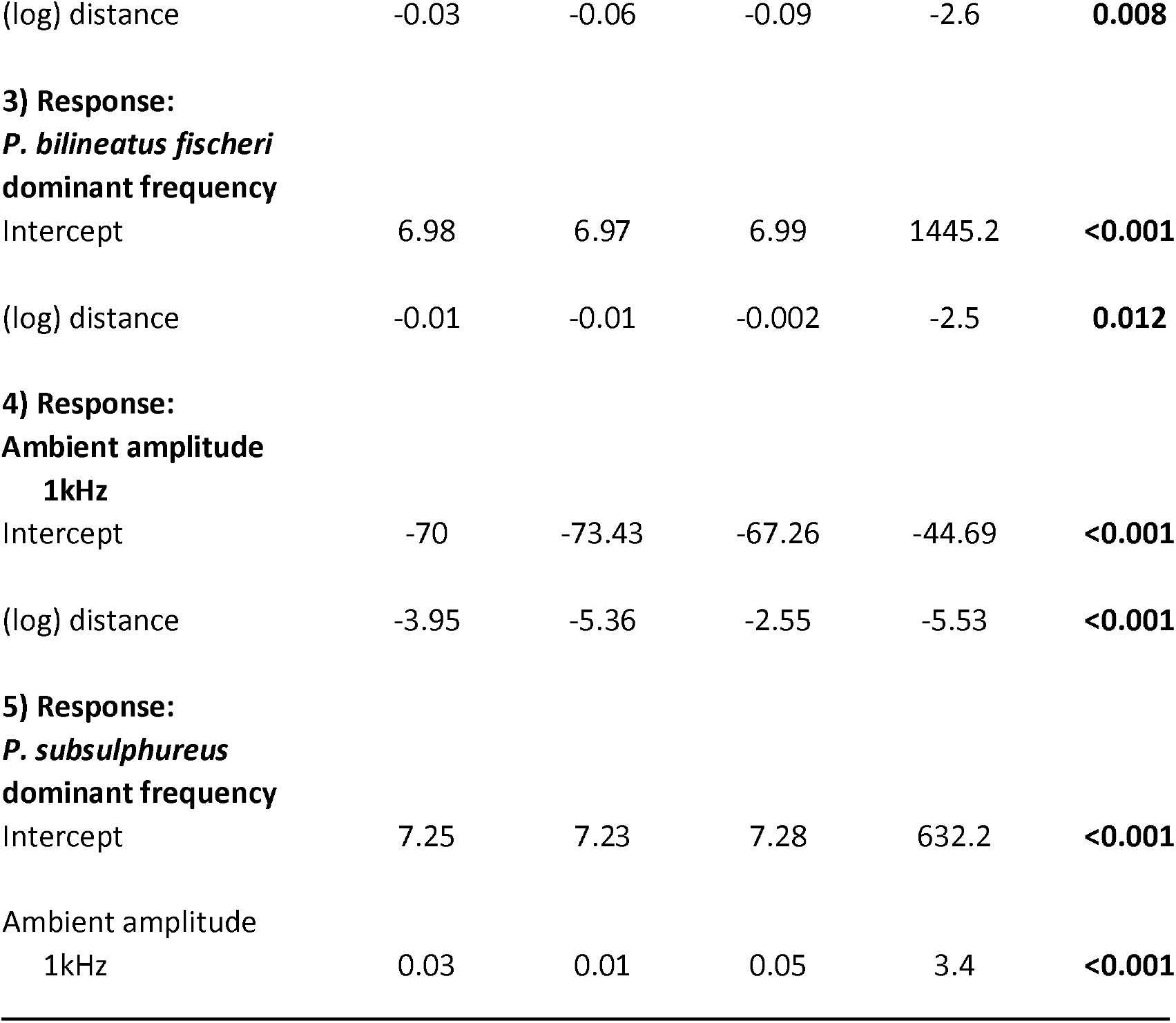
Best fit Gaussian generalized linear mixed models output showing the relationship between (log) dominant frequency and (log) distance from the coast for *subsulphureus* (AICc: −138.24) (1),*pusillus* (2), *fischeri* (3) as well as relationship between surf sound ambient noise and distance from the coast in Cameroon (4) and between *subsulphureus* dominant frequency and ocean surf sound (5). Estimates and their lower (LCL) and upper (UCL) confidence limits are presented.

**Fig. 2.**
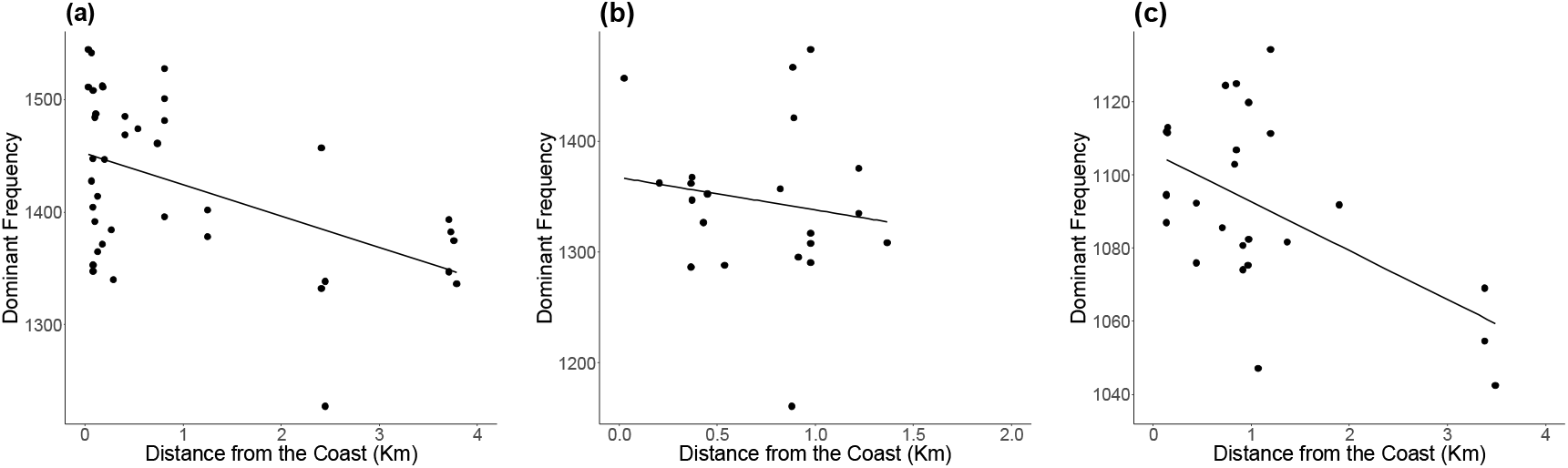
Plots showing the relationship between dominant frequency and distance from the coast in (a) *subsulphureus*, (b) *pusillus* and (c) *fischeri* before controlling for other possible effects.

Our results also show a strong significant decrease of low-frequency ambient noise (1kHz) with log-distance from the coast in Cameroon as well as a significant positive relationship between *subsulphureus* dominant frequency and ambient noise amplitude at 1kHz (Fig 3, Table 1).

**Fig. 3.**
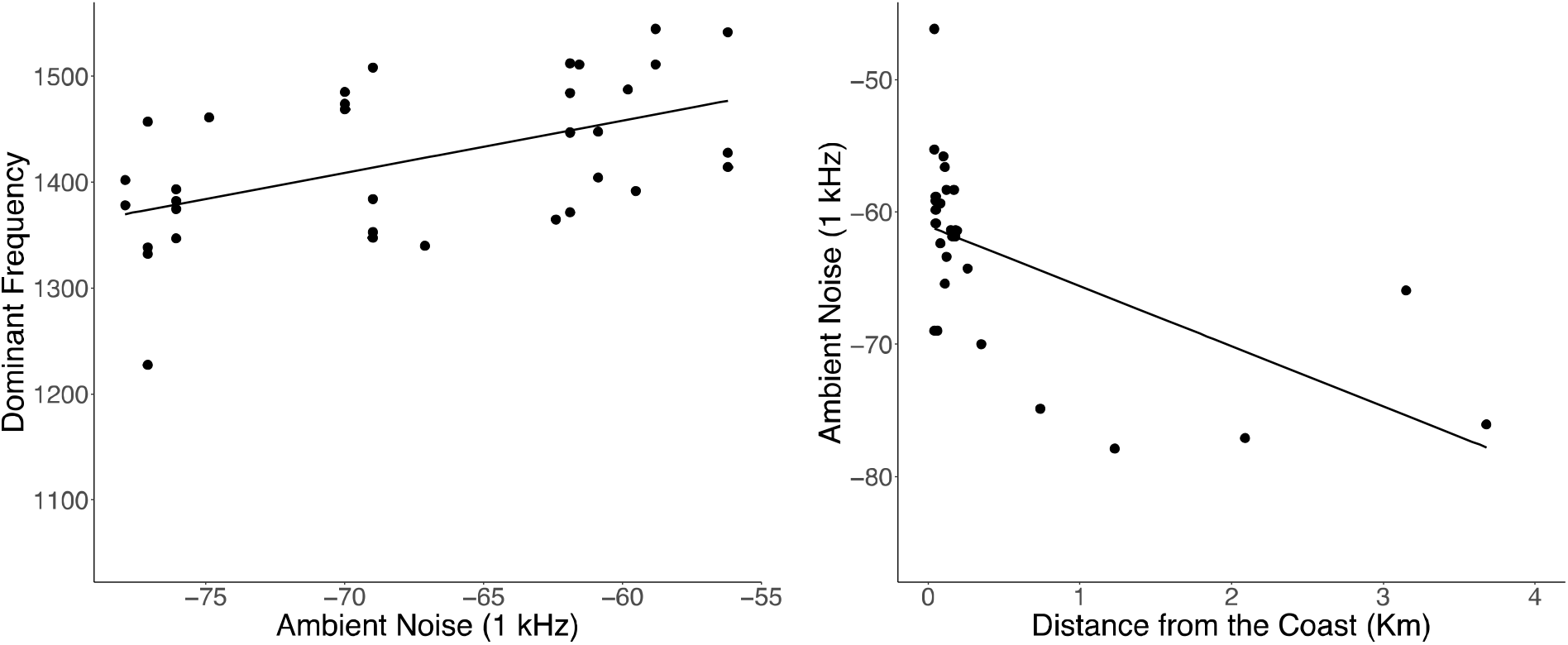
Plots showing (a) the relationship between *subsulphureus* dominant frequency and ambient noise and (b) between ambient noise and distance from the coast in Cameroon.

## Discussion

We have shown that tinkerbirds sing at higher frequencies the closer they are to the coast and as the amplitude of low-frequency ambient noise from ocean surf increases. Our results are in line with the hypothesis that, as in the case of anthropogenic noise, natural ambient noise affects acoustic signalling in birds (Kirschel et al. 2009a; Davidson et al. 2017). We show that the masking effect of a natural low-frequency noise can affect vocalizations of animals that lack the capacity to develop vocalisations through auditory feedback. Higher dominant frequency may confer an adaptive advantage to coastal populations of the three species tinkerbirds because they benefit from increased transmission of signals to intended receivers. Therefore, individuals with higher dominant frequency songs may have higher fitness at coastal sites. Low frequency natural noise such as from ocean surf, rivers and waterfalls can have a profound effect on auditory communication, as shown in concave-eared torrent frog (*Amolops tormotus*) whose calls include ultrasound elements in their preferred habitat alongside fast-flowing streams (Feng et al. 2006). Similar results have also been found in another study, where support for the acoustic adaptation hypothesis has been demonstrated when comparing torrent frogs to other species living in different habitats (Goutte et al. 2018). Tinkerbirds are not restricted to such noisy environments, yet divergence in frequency appears to occur in spite of ongoing gene flow with adjacent inland populations.

The pressure imposed by ocean surf low-frequency noise may have strong effects on how species interact acoustically because of potential interference with their vocalizations in the frequency domain. The effects of low-frequency ambient noise are likely to have a stronger effect on species vocalizing at lower frequency and especially in birds that lack vocal learning, such as tinkerbirds (Goodwin and Shriver 2011; Halfwerk et al. 2011). In this study, *fischeri* is the species with the lowest dominant frequency and therefore may be subjected to a greater pressure by ocean surf. In Kenya, it co-occurs with two other *Pogoniulus* tinkerbirds: *P. pusillus* and eastern green tinkerbird *P. simplex* (hereafter *simplex*), both of which sing at higher frequencies than *fischeri*. Indeed, *simplex* sings a trilled song not unlike that of *fischeri*. It is therefore possible that continental populations of *fischeri* are constrained to avoid the masking effect of low-frequency ocean surf sound by increasing their dominant frequency because an increased pitch would result in greater interference with the two competitors. Indeed, an increase in dominant frequency in continental populations of *fischeri* could lead to song overlap in the frequency domain with its two congeners (Fig. 4b). Stabilising selection might maintain *fischeri* song frequency at a level that best reduces the masking effects of surf sound while maintaining sufficient frequency differences between *fischeri* and other tinkerbird species. Coastal *fischeri* sing a much faster trilled song than other forms of *P. bilineatus* (Nwankwo et al. 2018) and the rapid repetition of pulses might itself be an adaptation to its sound environment in coastal forests. An alternative hypothesis is that *fischeri* song might have evolved by convergent character displacement to facilitate interspecific territoriality with *simplex* (e.g., Kirschel et al. 2019). The observed increase of frequency in *fischeri* might also reflect the increase in dominant frequency in *pusillus* song, as its frequency range may depend on *pusillus* minimum frequency. Hence, the observed decreasing pattern in *fischeri* dominant frequency with distance from the coast may in part be an effect of variation in *pusillus* song with distance from the shore. A similar, if not stronger, correlation between frequency ranges is expected to occur between *fischeri* and *simplex*, given the similarity of the song between the two species. However, we did not have access to a suitable sample of *simplex* recordings to test this hypothesis. Further work is needed to investigate the extent to which *fischeri* song may also vary because of interactions with its congeners.

**Fig. 4.**
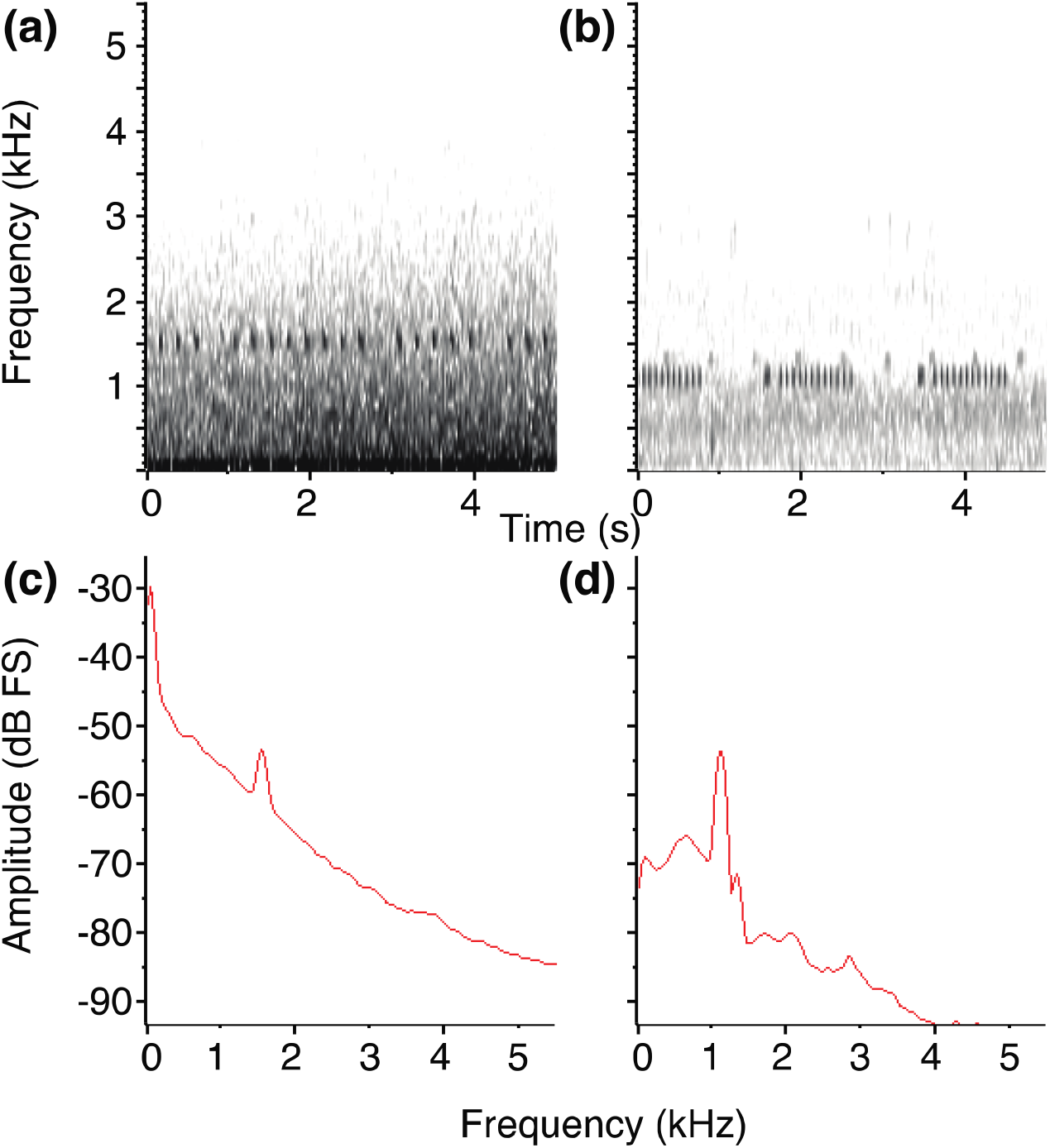
Figure showing the masking effect of ocean surf sound in spectrograms (a-b) and power spectra profiles (c-d) of *subsulphureus* (left panels), *pusillus* and *fischeri* (right panels) vocalizations, with *subsulphureus* song being recoded at 68 m form the shore and *pusillus* and *fischeri* recording at 330 m.

Higher dominant frequency have been suggested to be a consequence of an increased song amplitude in non-passerines (Elemans et al. 2008; Nemeth and Brumm 2010; Nemeth et al. 2012), whereas this is not always the case in passerines, which present higher vocal flexibility (Zollinger et al. 2017). An increased amplitude can be an adaptation to noisy environments according to the Lombard effect, which occurs when frequency range of the vocalizing animal and the background noise overlap (Brumm and Todt 2002). In our study, ocean surf sound widely overlaps with tinkerbirds song frequencies (Fig. 4), therefore one possibility is that increased dominant frequency in tinkerbird song at coastal sites is a consequence of raised vocal amplitude. The Lombard effect is a common trait in many bird clades including passerines (Brumm and Todt 2002), Galliformes (Brumm et al. 2009) and even in Paleognathae species such as tinamous (Schuster et al. 2012). The ancestral nature of the Lombard effect suggests it occurs independently of the ontogeny of vocal learning in birds (Brumm et al. 2009; Brumm and Zollinger 2011) and increased frequencies in tinkerbird song might also be a consequence of increased vocal amplitude. This phenomenon has been observed in other birds that lack song learning (Schuster et al. 2012). However, we did not specifically test whether the increased dominant frequency occurs as a consequence of the Lombard effect in tinkerbirds, but our results highlight this as a compelling area for future investigations.

Singing higher pitch songs in coastal sites may be an advantage in tinkerbirds, as higher frequency songs often represent a selected trait by females (Hasegawa and Arai 2016). Also, an increased pitch may result in an increased detectability by opposite-sex individuals. Assuming that song frequency is correlated with amplitude, increased frequency would result in a far-reaching signal which may further aid mate attraction. Similarly, in territorial contests, higher pitch song may result in a larger active space (Brumm and Todt 2002) – a potential advantage in territorial birds like tinkerbirds. However, pitch has been shown in many birds to be negatively correlated with body size (Ryan and Brenowitz 1985; Brumm and Goymann 2017, Kirschel et al. 2020, *in press*), whereas it does not seem to affect song amplitude (Brumm 2009). Hence, any relative advantage in terms of signal transmission may be counterbalanced by increased aggression from larger males, as higher frequency song may be interpreted as a sign of weakness (Kirschel et al. 2020, *in press*). Ocean surf sound is a continuous noise which pressure acts over evolutionary timescales on birdsong, therefore the trade-offs between the potential advantages of increased mate attraction and at the same time increased territorial response from other males may have had profound evolutionary implications in shaping tinkerbird acoustic signals.

In this paper, we show that three tinkerbird species sing at a higher dominant frequency the closer they are to the coastline. We suggest that low-frequency noise from ocean surf imposes a selective pressure on tinkerbird acoustic signalling, and higher dominant frequency songs may be selected because they reduce the masking effect of ocean surf sound. This effect might be boosted if an increase in dominant frequency is accompanied by an increase in amplitude. We predict that an increase in dominant frequency will occur but caution that overlapping frequencies with related species might influence acoustic competition, as might occur in *fischeri* where it coexists with *pusillus* and *simplex*. Our results show that natural ambient noise has a similar impact to anthropogenic noise even on birds that do not learn their songs, in line with the effects of natural ambient noise on oscine passerine vocalizations (Davidson et al. 2017). We believe that our results can be extrapolated in other contexts of background noise, including anthropogenic noise pollution, and therefore represent a baseline for further studies on the effect of background noise on bird song.

## Notes

### Competing Interest Statement

The authors have declared no competing interest.

