## Supplementary material for "Ambient noise from ocean surf drives frequency shifts in non-passerine bird song": Table S1

**Table S1** Full Gaussian generalized linear mixed model output (AICc: -122.87) showing the relationship between *subsulphureus* dominant frequency, (log) distance and ambient noise measured at 1 kHz. Estimates and their lower (LCL) and upper (UCL) confidence limits are presented.

|  |  | **Estimate** | **LCL** | **UCL** | ***z*** | ***p*** |
| --- | --- | --- | --- | --- | --- | --- |
| ***P. subsulphureus*** | |  |  |  |  |  |
| **dominant frequency** | |  |  |  |  |  |
| Intercept |  | 7.26 | 7.23 | 7.29 | 474.4 | **<0.001** |
| (log) distance | | 0.001 | -0.014 | 0.016 | 0.2 | 0.872 |
| Ambient amplitude | | 0.033 | 0.005 | 0.062 | 2.3 | **0.02** |
| 1kHz |  |  |  |  |  |  |
